# A potential genomic recombination site upstream of the *rfb* locus in *Leptospira interrogans* is associated with serogroup Serjoe and serovar Hardjo classification

**DOI:** 10.1101/771931

**Authors:** Maria Raquel V. Cosate, Tetsu Sakamoto, Élvio C. Moreira, José Miguel Ortega, Rômulo C. Leite, João Paulo Haddad, Tiago Antônio de Oliveira Mendes

## Abstract

Leptospirosis is a zoonotic disease caused by pathogenic spirochetes of the genus *Leptospira*. It has a global distribution and affects domestic animals, including cattle. In livestock production, *Leptospira interrogans* serogroup Sejroe serovar Hardjo is the major reproductive disease leading to economic losses. The whole-genome sequence of the first Brazilian clinical isolate classified as *L. interrogans* serogroup Sejroe serovar Hardjo strain Norma enabled the evaluation of its genomic features. Here, we investigated particularities of this isolate, obtained from a leptospirosis outbreak. Bioinformatic analysis using the *L. interrogans* serovar Hardjo str. Norma was applied as a reference for genomic evaluation and comparative analysis among *L. interrogans* and *L. borgpetersenii* serovars. Our data suggest the occurrence of genomic recombination in *L. interrogans* serovar Hardjo str. Norma encompassing 45 Kb located upstream of the *rfb* locus. A hallmark of genetic evolution was predicted through an orthologue analysis that identified that sugar enzymes associated with carbohydrate and lipid biosynthesis and metabolism composed this genetic module. Comparative genomics revealed a wide range of relatedness among the bacterial strains of serogroup Sejroe that are classified as *L. interrogans* and *L. borgpetersenii* species. Furthermore, identification of an *IS3* family suggests a genetic recombination site in *L. interrogans* serovar Hardjo str. Norma that is distinct among *L. interrogans* serovars and may contribute to clarify the taxonomic classification of *Leptospira* spp.

**Impact Statement:** Leptospirosis remains an important neglected disease with worldwide distribution. This zoonotic disease impacts in the livestock production and the bovine infection is currently associated to species *L. borgpetersenii* and *L. interrogans* serovar Hardjo. *L.interrogans* serovar Hardjo infection is recognized as reproductive disease associated with abortion and economic lost. In this context, we studied a unique whole genome sequence of *L. interrogans* serovar Hardjo subtype Hardjo-prajitno isolated from bovine leptospirosis outbreaks in Brazilian dairy farm, one of the greatest country of milk production in world. We compared *L. interrogans* and *L. borgpetersenii* genomes with *L. interrogans* serogroup Sejroe serovar Hardjo subtype Hardjo-prajitno focusing on *rfb* locus and sugars biosynthesis. Leptospira spp. taxonomy and serology information are strictly associated with rfb locus and we found high correlation among bacterial strains classified in serogroup Sejroe. Although *L. interrogans* and *L. borgpetersenii* classified in serogroup Sejroe possess a greater genetic correlation, we uniquely identified that serovar Hardjo strains often possess identical loci carrying predicted sugar biossinthesis genes and mobile elements. The *Sru* (**S**ejroe specific **R**fb **U**pstream locus) locus associated to *rfb* locus probably contribute to *Leptospira* spp. genetic information concerning serogroup and serovar degrees of taxonomic and serology in this microbiology field.

**Data Summary:** All *Leptospira* spp. genome sequences used in this study were retrieved from National Center for Biotechnology Information (NCBI) (Table 1) with NCBI ID: NZ_CP006723.1, NZ_CP012603.1, NC_004342.2, NZ_CP011934.1, NZ_AKXA02000040.1, NZ_CP013147.1, NZ_CP012029.1, NZ_CP015048.1, NC_008508.1, GCA_000216175.3,GCA_000244115.3, NC005823.1, GCA_000244395.3, GCA_000346975.1.

**Highlights:**

1. The study identified potential molecular features associated with serovar Hardjo subtype Hardjo-prajitno, present in both *L. interrogans* and *L. borgpetersenii*, by comparative genomics.
2. A new potential recombinant site found upstream of the *rfb* locus contains proteins that correlate with serogroup taxonomy in the *Leptospira* genus.
3. The proteins encoded in the recombinant locus are associated with the synthesis of serological surface determinants such as carbohydrates and lipopolysaccharides.

## Introduction

Leptospirosis is an important zoonosis with a worldwide distribution and is caused by bacteria classified in the *Leptospira* genus (1). This genus includes pathogenic and saprophytic species, in which *Leptospira interrogans* is the most prevalent species that is directly associated with human and animal disease (2, 3). Currently, there are approximately 22 *Leptospira* species distributed in more than 300 serovars (4). Previous studies have described that serovar and serogroup classifications are directly associated with the biosynthesis and composition of lipopolysaccharides and have no correspondence with taxonomic classification (1, 5, 6). Therefore, different *Leptospira* species can be classified in the same serogroup and serovar, as in serogroup Sejroe serovar Hardjo; for example, the *Leptospira* species *L. interrogans* and *L. borgpetersenii* belong to this group (1).

Although *L. interrogans* serogroup Sejroe serovar Hardjo subtype Hardjoprajitno is highly pathogenic for cattle, resulting in low milk production and increased abortion frequency, *L. borgpetersenii* serogroup Sejroe serovar Hardjo subtype Hardjobovis is more adapted to bovine species, with reduced clinical signs and pathogenesis (7,8,9). Differences among those bacterial subtypes include differences in geographical regions of isolation, genome composition, capacity for environmental survival and the lipopolysaccharide composition associated with the *rfb* locus (9, 10, 5, 6, 11). In *L. borgpetersenii* and *L. interrogans* of the same Hardjo serovar, the *rfb* loci is composed of 31 and 32 open read frames (ORFs), respectively. The additional ORF in *L. interrogans* is located between ORF4 and ORF5 (6, 12). Most *rfb* ORFs are associated with the biosynthesis of sugars, such as rhamnose, mannosamine, fucosamine and galactose (5,6,11). Previous analyses of the *rfb* locus in the *L. borgpetersenii* serovar Hardjo genome have also identified a set of mobile elements classified as insertion sequence (IS) family 3 in the intergenic region between ORF14 and ORF15; this insertion sequence is supposedly absent in the genome of *L. interrogans* serovar Hardjo subtype Hardjoprajitno strain Hardjoprajitno (6). Molecular analyses based on PCR and Sanger sequencing of specific target sequences showed that the region spanning ORF1 to ORF15 of the *L. interrogans* Hardjoprajitno *rfb* locus has high similarity with *L. borgpetersenii* serovar Hardjo, while the region spanning from ORF15 to ORF31 has high similarity with *L. interrogans* serovar Copenhageni (11). Furthermore, phylogenomics and comparative genomic analyses of the two *L. interrogans* genome samples from serovar Hardjo and 20 genomes from other serovars identified differences in transporters associated with drug resistance, PIN domain evolution and chromosome features, including the *rfb* locus (13). Although molecular characterization of *Leptospira* spp. has focused on the *rfb* locus, genomic characteristics that are probably associated with the Hardjo phenotype and potential evolutionary mechanisms associated with the distribution of this serovar in both species (*L. interrogans* and *L. borgpetersenii*) have not been considered.

The recent publication of two *L. interrogans* serogroup Sejroe serovar Hardjo subtype Hardjoprajitno genomes and the availability of genomes from *L. borgpetersenii* serovar Hardjo bovis enable a more accurate comparison to identify genomic features associated with the phenotype and evolution of this serogroup (13). Here, we performed a comparative analysis among Hardjo and non-Hardjo strains. Our findings suggest that the Hardjo phenotype is not completely delimited to the rfb locus but includes the *sru* locus (**S**ejroe-specific ***Rfb* U**pstream locus) composed of 43 genes located upstream from the rfb locus. An orthologous analysis suggests that this region is enriched with genes associated with sugar biosynthesis and metabolism. The identification of the mobile element IS3 inside this region suggests a hot spot recombination site that is associated with the horizontal transfer of Hardjo molecular features among *L. interrogans* and *L. borgpetersenii*.

## Methods

### *Leptospira* spp. genome sequences

A total of nine *Leptospira* spp. genome sequences were selected and retrieved from the National Center for Biotechnology Information (NCBI) public database. For this study, we selected *Leptospira* strains that were considered to be reference entries and those classified in the serogroup Sejroe, which includes entries from *L. interrogans* and *L. borgpetersenii* species (Table 1). Nucleotide and predicted amino acid sequences were compared using *L. interrogans* serogroup Sejroe serovar Hardjo subtype Hardjoprajitno str. Norma as reference. Additional database searches were performed using BLASTn and BLASTp, available from NCBI (https://www.ncbi.nlm.nih.gov/).

**Table 1:** *Leptospira* spp. strains used in this study.

| Species | Serogroup | Serovar | Strains | NCBI ID |
| --- | --- | --- | --- | --- |
| *L. interrogans* | Icterohaemorrhagiae | Lai | 56601 | PRJNA293 |
|  | Icterohaemorrhagiae | Copenhageni | L1-130 | PRJNA10687 |
|  | Sejroe | Hardjo | Norma | PRJNA185511 |
|  | Sejroe | Hebdomadis | C401 | FJ976889 |
|  | Sejroe | - | Brem 329 | PRJNA167229 |
|  | Sejroe | Hardjo | Hardjoprajitno | PRJNA296687 |
|  | Icterohaemorrhagiae | Canicola | LT-1962 | PRJNA65063 |
| L. borgpetersenii | Sejroe | Hardjo-bovis | L550 | PRJNA16146 |
|  | Sejroe | Hardjo-bovis | BK-30 | PRJNA296679 |

### Genome comparative analysis

We compared the genome sequence of *L. interrogans* serovar Hardjo str. Norma with eight *Leptospira* isolates that had publicly available genomes in the NCBI and PATRIC databases (14). Initially, the Proteome Comparison Tool implemented in PATRIC (https://www.patricbrc.org/) was used to generate a matrix of the bidirectional best hits between the proteins from all genomes using the BLASTp algorithm (15). For each best hit, we recovered information about the coverage, sequence identity, protein annotation and genomic coordinates (Supplementar Table 1). The best hit pairs were used to build syntenic maps.

### Gene ontology analysis

The protein sequences were used as input in the BLASTp tool implemented in the UniProt database (release 2019_02). The best hit was selected for each protein, and the descriptive gene ontology (GO) terms regarding biological processes and molecular functions were retrieved from each best hit. The GO terms were grouped into 15 general categories (transferase activity, oxidoreductase activity, carbohydrate biosynthesis, ligase activity, transaminase activity, ion binding, isomerase, lipopolysaccharide biosynthesis, ATPase activity, phosphatidylinositol phosphorylation and NAD binding).

### Molecular phylogenetic analysis

To perform phylogenetic studies, included the sequence information from the isolates *L. interrogans* serovars Medanensis (NCBI ID: PRJNA74127), Manilae (NCBI ID: PRJEB23342), and Linhai (NCBI ID: PRJNA217894) and *L. borgpetersenii* serovars Hardjo-bovis (NCBI ID: PRJNA296679) and Ballum (NCBI ID: PRJNA255706) to along with the selected genomes due to epidemiological correlations with *L. interrogans* serovar Hardjo subtype Hardjo-prajitno (Supplementar Table 6). Putative homologous sequences of each genomic region of interest in the other *Leptospira* spp. were retrieved from the GenBank database through a BLAST search (15). This search was performed on the NCBI web server (http://blast.ncbi.nlm.nih.gov/Blast.cgi) by submitting the nucleotide and translated sequences obtained from our isolates as queries against the refseq_genomic and UniProt databases, respectively. The sequence selection was performed based on the best hits with at least 40% sequence coverage and 50% identity. Selected sequences obtained from the database were aligned using MUSCLE (16)]. All phylogenetic trees generated in this work were inferred by the maximum likelihood method with model maximum composite likelihood and 1,000 bootstrap replications, implemented in MEGA 6 (17).

## Results

### The *rfb* locus from *L. interrogans* serovar Hardjo is a hybrid between Hebdomadis and Hardjo-bovis

The r*fb* locus has been used to classify *Leptospira* into serogroups (1, 18, 19). Thus, we selected *Leptospira* species that are historically related with serogroup Sejroe and serovar Hardjo (Table 1) to evaluate the *rfb* locus conservation between different *Leptospira* species. The *rfb* locus of *L. interrogans* serogroup Sejroe serovar Hardjo subtype Hardjoprajitno strain Norma has 37 Kb and is composed of 32 predicted ORFs (Figure 1A). Comparative analysis showed that the Norma strain has identical gene synteny with *L. interrogans* str. Brem 329, *L. interrogans* serovar Hardjo str. Hardjoprajitno and *L. borgpetersenii* serovar Hardjo-bovis L550 (Figure 1A).

**Figure 1:**
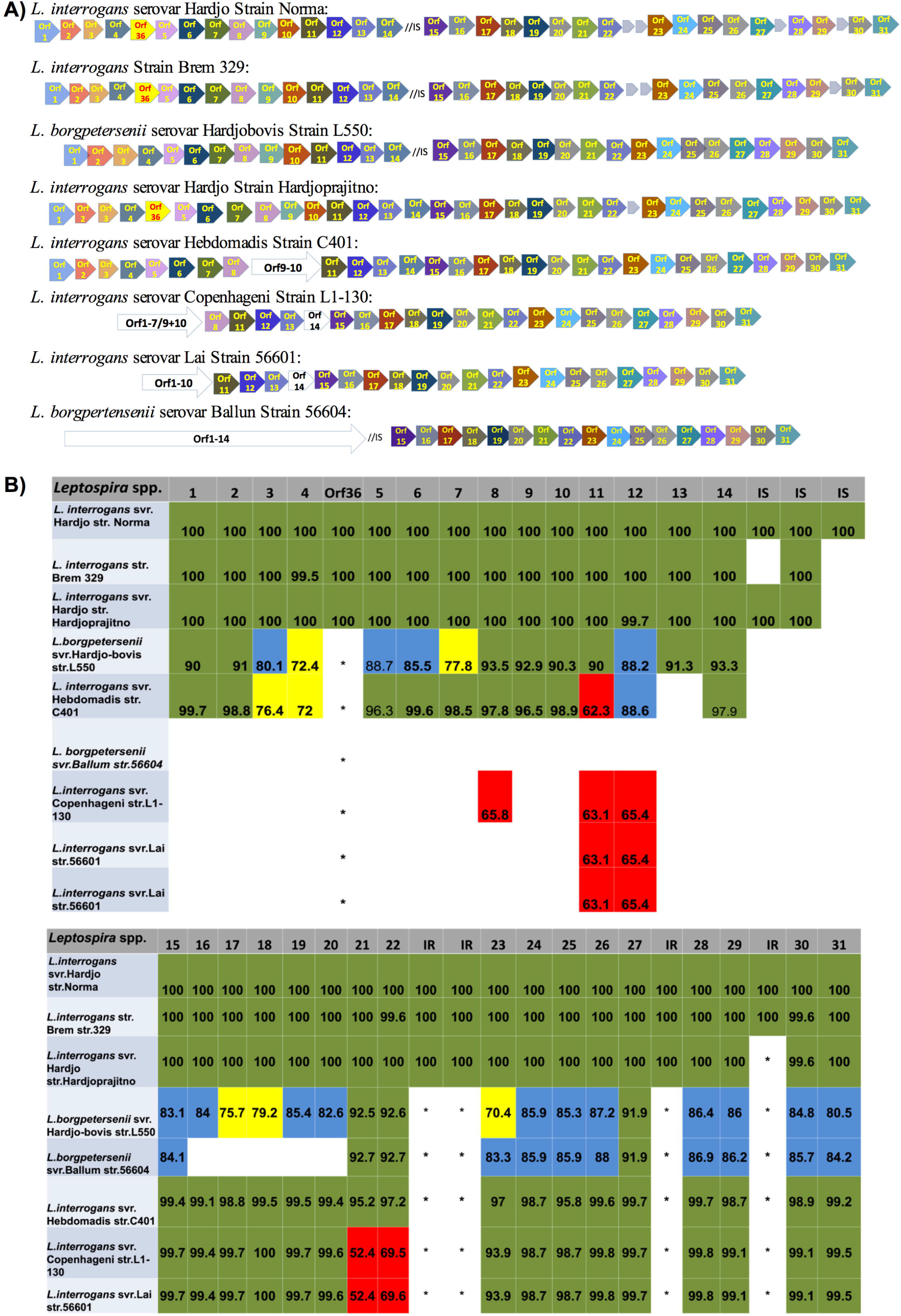
The *rfb*-locus synteny of *L. interrogans* and *L. borgpetersenii* serovars. (A) Comparative organization of *rfb* locus among *Leptospira* spp. Insertion sequence (//IS) is shown between Orf14 and Orf15. Intergenic sequences are located between Orf22-Orf23, Orf27-Orf28 and Orf29-Orf30. *Leptospira* strains are organized according to the similarity with str. Norma. *L.interrogans* str. Brem, *L.borgpetersenii* serovar Hardjo-bovis str.L550, *L.interrogans* serovar Hardjo str.Hardjoprajitno, *L.interrogans* serovar Hebdomadis str.C401, *L.interrogans* serovar Copenhageni str.L1-130 and L.interrogans serovar Lai str.56601. The white arrows represents lower identities (<40%) among *L.interrogans* serovar Copenhageni str. L1-130, *L.interrogans* serovar Hebdomadis str.C401 and *L.interrogans* serovar Lai str.56601 with the reference *L.interrogans* serovar Hardjo str. Norma. (B) ***rfb* locus identity among *Leptospira* spp.** Nucleotide identities of *rfb* locus among leptospira serovars. Green (90%-100%), Blue (80% - 89%), Yellow (70%-79%), Red (<70%) and white represent low identities (<50%) or abscense.

Previous work indicates that in the *L. interrogans* serovar Hardjo subtype Hardjoprajitno *rfb* locus, ORF1 to ORF14 were probably acquired from *L. borgpetersenii* serovar Hardjo-bovis and ORF15 to ORF31 were acquired from *L. interrogans* serovar Copenhageni L1-130 (20). Here, we observed that ORF11 and ORF13 are highly conserved among *L. interrogans* and *L. borgpetersenii* serovar Hardjo when compared among serogroup Sejroe bacterial strains. We also detected a mobile insertion sequence (IS) element between ORF14 and ORF15 from *L. interrogans* serovar Hardjo strain Norma, Brem 329 and *L. borgpetersenii* (Figure 1A). Furthermore, we did not identify those elements in *L. interrogans* serovar Hardjo strain Hardjoprajitino, *L. interrogans* serovar Hebdomadis strain C401, *L. interrogans* serovar Copenhageni or *L. interrogans* serovar Lai.

Although previous studies of the *rfb* locus in *L. interrogans* serovar Hardjo described a high similarity from ORF15 to ORF31 with *L. interrogans* serovar Copenhageni, here we observed the highest identities of nucleotide sequences with *L. interrogans* serovar Hebdomadis, which is also classified in serogroup Sejroe, and included ORF21 (95.2% and 92.5% with Hebdomadis and Hardjo-bovis, respectively) and ORF22 (97.2% with Hebdomadis and 92.6% with Hardjo-bovis) (Figure 1B and Supplementar Table 2). These areas were previously described as being more similar to serovar Hardjobovis (11, 6). In contrast, the analysis of *L. interrogans* serovar Copenhageni showed identities of 52.4% and 69.5% for those two genes (ORF21 and ORF22), respectively.

Despite the high similarity of the *rfb* locus from *L. interrogans* serovar Hardjo with *L. interrogans* serovar Hebdomadis and *L. borgpetersenii* serovar Hardjo-bovis, we observed a unique nucleotide sequence present in the Hardjoprajitino genome (Figure 1B, Supplementar Table 2). Although we did not identify *L. interrogans* str. Brem 329 in the database as serovar Hardjo, here we are considering this classification due to the genome conservation within *L. interrogans* serovar Hardjo. ORF36 was detected only in *L. interrogans* serovar Hardjo. Prediction sequence analysis showed a high identity with 2-oxoglutarate (2OG)-FeII (dependent oxygenase), a class of enzymes that are widespread in bacteria and are associated with DNA and mRNA repair and synthesis (21). In addition, we identified four hypothetical proteins located between ORF22-ORF23, ORF27-ORF28 and ORF29-ORF30, where strain Hardjo-prajitno lacks the latter one (Figure 1B).

### The region upstream of the *rfb* locus in *L. interrogans* strain Hardjo and *L. borgpetersenii* serovar Hardjobovis has conserved synteny

Since the beginning of the *rfb* locus from serovar Hardjo is more closely related to *Leptospira* strains that belong to *L. borgpetersenii* serovar Hardjobovis, we evaluated whether the region upstream of the *rfb* locus is maintained. The 45.1 Kb genome segment upstream of the *rfb* locus is composed of 43 ORFs with identical synteny to *L. borgpetersenii* serovar Hardjo-bovis (Figure 2). This genome region is referred to as the *sru* locus. The synteny was also conserved in other leptospiras from serogroup Serjoe and serovar Hardjo, including *L. interrogans* serovar Hardjo str. Hardjo-prajitno and *L. interrogans* str. Brem 329. On the other hand, *L. interrogans* serovar Hebdomadis showed an absence of ORF13 to ORF16; in its place, we identified a mobile element classified as an insertion sequence (IS) (Figure 2 and Supplementar Table 3). In contrast, the region upstream of the *rfb* locus in serovars Lai and Copenhageni showed very different gene organization in comparison with the other *Leptospira* strains in the analysis. Positive matches were observed only for ORF30, Orf33, Orf34, Orf35, Orf36, Orf41 and Orf43 (Figure 2 and Supplementar Table 3). The functional annotation of genes comprising the *sru* locus showed that most coded for enzymes involved in transferase activity, including methyltransferases, cytidylyltransferase and nucleotidyltransferase (Figure 3, Supplementar Table 4). Interestingly, the second expressive functional group had proteins associated with sugar metabolism pathways, such as oxidoreductase, ligase, transaminase and ion binding activity (Figure 3). The predicted protein annotation included 6-phosphogluconolactonase (WP_060589238.1), phosphoheptose isomerase (WP_001047905.1) (GDP-L-fucose synthetase and N-acetylneuraminate synthase (Supplementar Table 4). We also identified 3-deoxy-manno-octusolonate cytidylyltransferase (WP_000848241.1) and galactoside O-acetyltransferase (WP_002189008.1), which are associated with lipopolysaccharide biosynthesis. Thus, ORFs 13, 14, 15, 16, 37 and 40 have a high potential of being correlated with serovar Hardjo.

**Figure 2:**
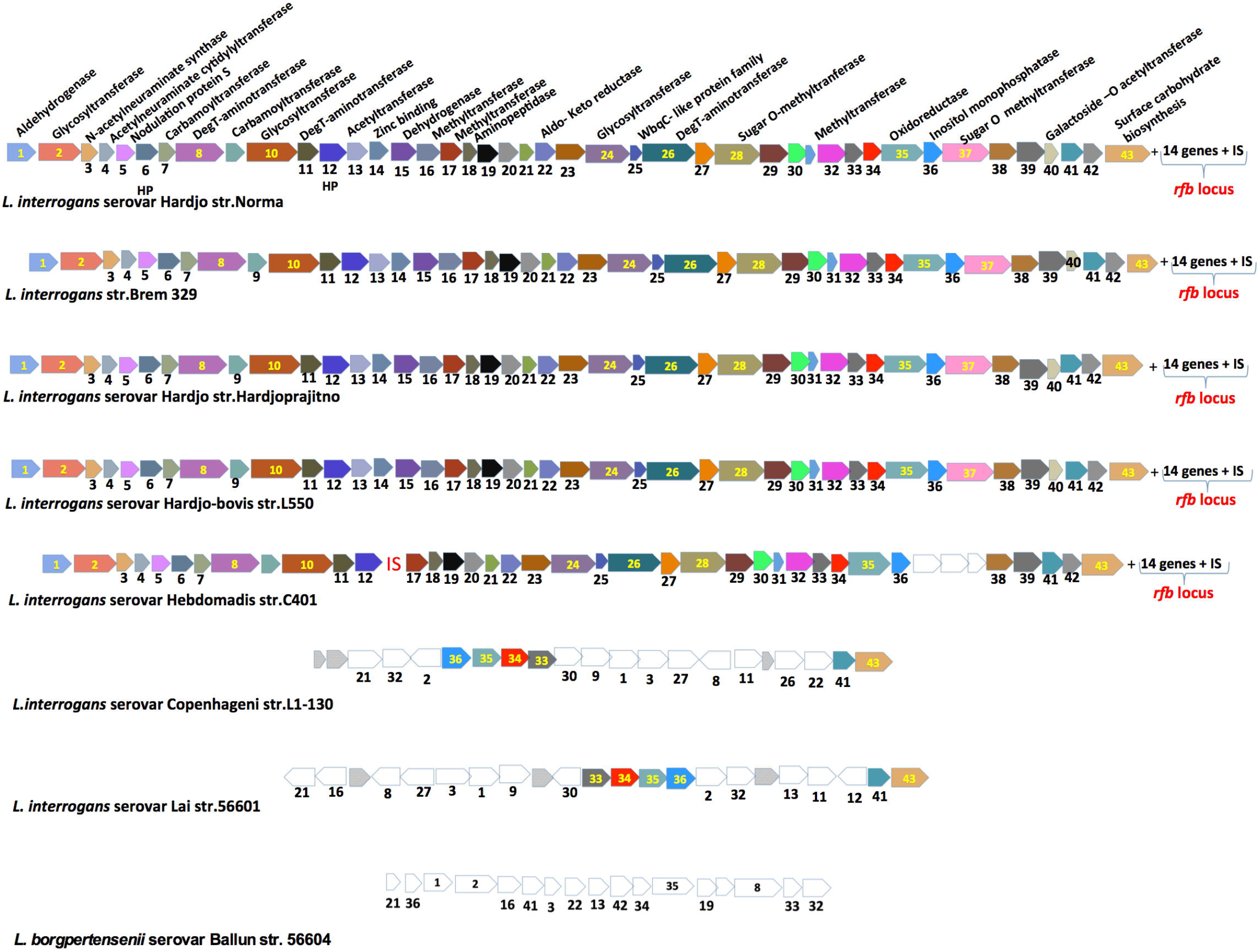
The structure of *Sru* locus in *Leptospira spp*. Schematic organization of the genes in the Sru locus of *L.interrogans* serovar Hardjo str.Norma. A total of 43 Sru genes followed by 14 genes from rfb locus and the mobile element (IS) are showed. *Leptospira* strains are organized according to synteny similarity with *L.interrogans* serovar Hardjo str. Norma. Insertion sequence (IS) was identified in *L.interrogans* serovar Hebdomadis str.c401. The white arrows respresent low aligment matches (<40 of identity and <40% of coverage sequence).

**Figure 3:**
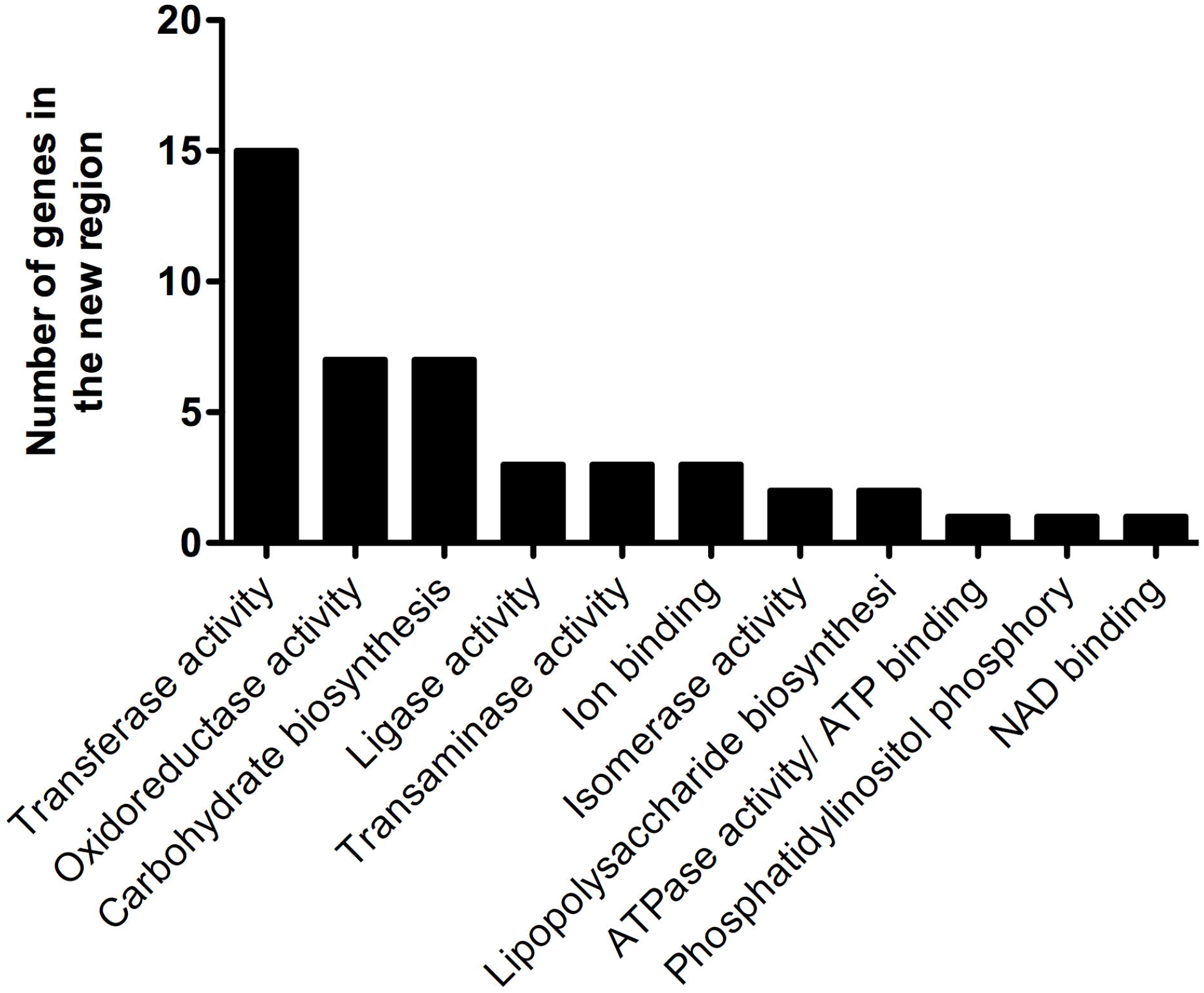
Functional analysis of genes identified in the *Sru* locus from *L. interrogans* serovar Hardjo. Gene ontology terms were identified for all genes predicted in the *L.interrogans* serovar Hardjo *Sru* locus. The terms were organized in descending order of frequency.

### Proteins in the *sru* locus show phylogenetic correlation with serogroup Sejroe serovar Hardjo

Because proteins in the *sru* locus showed a high conservation between serovar Hardjo subtype Hardjoprajitino and serovar Hardjo-bovis, we applied phylogenetic analysis to study the protein and nucleotide sequence evolution for each ORF in this region. After phylogenetic reconstruction for each gene marker, we observed whether the topology of the produced trees was in agreement with one of the taxonomic classifications: by species, serogroup, serovar or genotype (Figure 4 and Supplementary Figures 1-2). A total of 3 out of 43 nucleotide sequences from the *sru* locus resulted in phylogenetic models with cladistic separation between *Leptospira* species (Figure 5, Supplementar Figure 3). These genes are annotated as 2,4-dihydroxyhept-2-ene-1,7-dioic acid aldolase, inositol monophosphatase and Shikimate/quinate 5-dehydrogenase. Interestingly, a total of 22 genes (51,5% of the *sru* locus) showed phylogenetic trees that grouped serogroups within the same clade (Figure 5, Supplementar Figure 1). Moreover, four genes were found only in strains from serovar Hardjo (zinc binding, dehydrogenase, SAM-dependent methyltransferase, S-adenosylmethionine-dependent methyltransferase), and one gene was identified only in strains from the genotype Hardjoprajitino (galactoside O-acetyltransferase).

**Figure 4:**
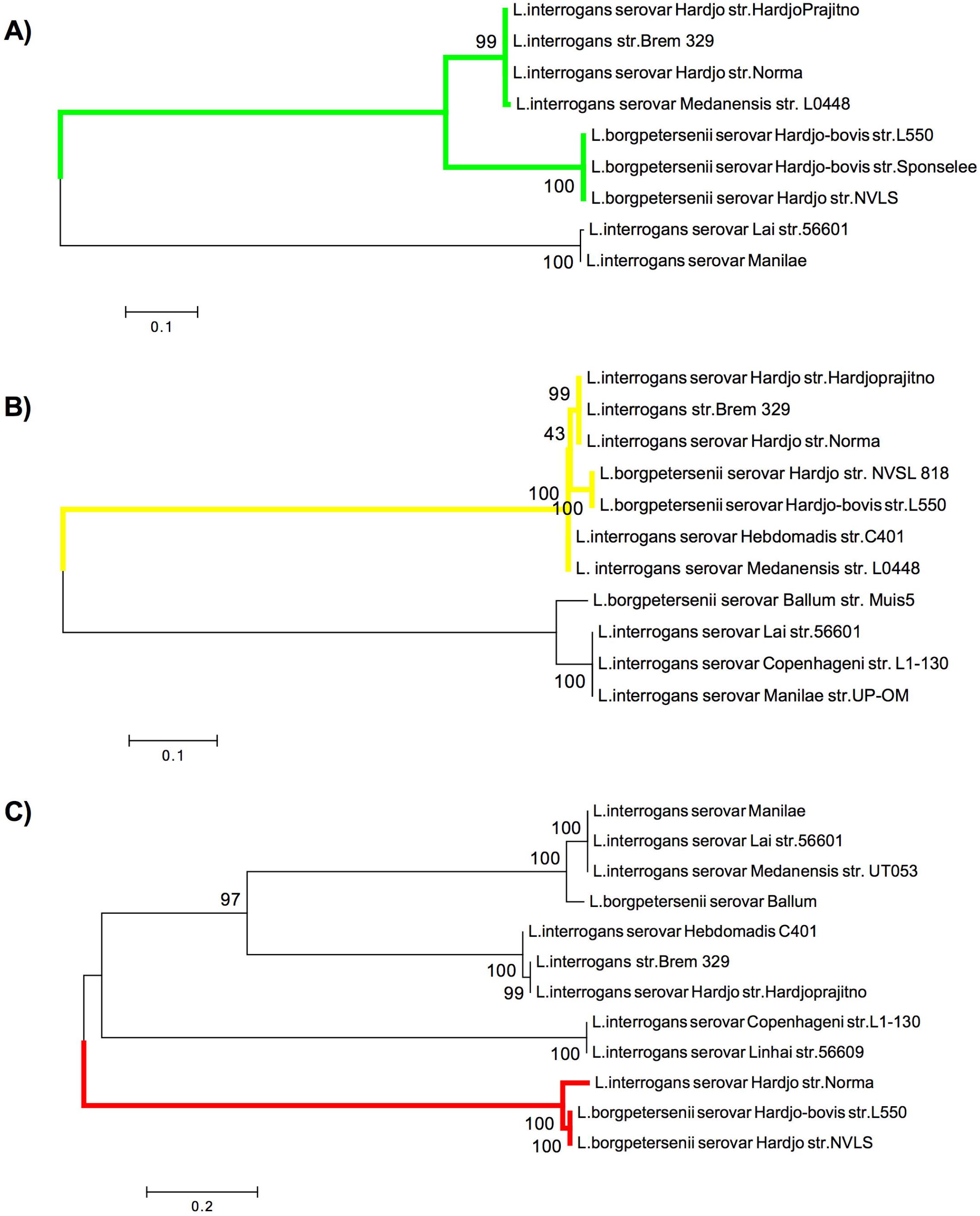
The phylogenetic trees of Sru genes from *Leptospira spp*. **(A**) Phylogeny based on the gene 2, 4 dihydroxyhept-2-ene-1,7-dioic acid aldolase. *L. interrogans* serovar Hardjo is clustered together diferent serovars classified in *L. interrogans* specie (green branches). *L.borgpetersenii* strains are shown in a distinct cluster. **(B)** Phylogeny based on the gene mehtyltransferase. Leptospira strains classified in serogroup Sejroe are clustered together (*L.interrogans* and *L.borgpetersenii* species) represented by yellow branches. L.interrogans serogroup icterohaemorrhagiae are clustered together. **(C)** Phylogeny based on the gene 2 DegT aminotransferase of *L. interrogans* are separeted into four clusters and *L.interrogans* serovar Hardjo str. Norma is clustered together with *L.borgeptersenii* serovar Hardjo-bovis str. L550 and NVLS (red branches). The phylogenetic trees were inferred by maximum likelihood method with 1000 bootstrap and Tamura Nei-Model.

**Figure 5:**
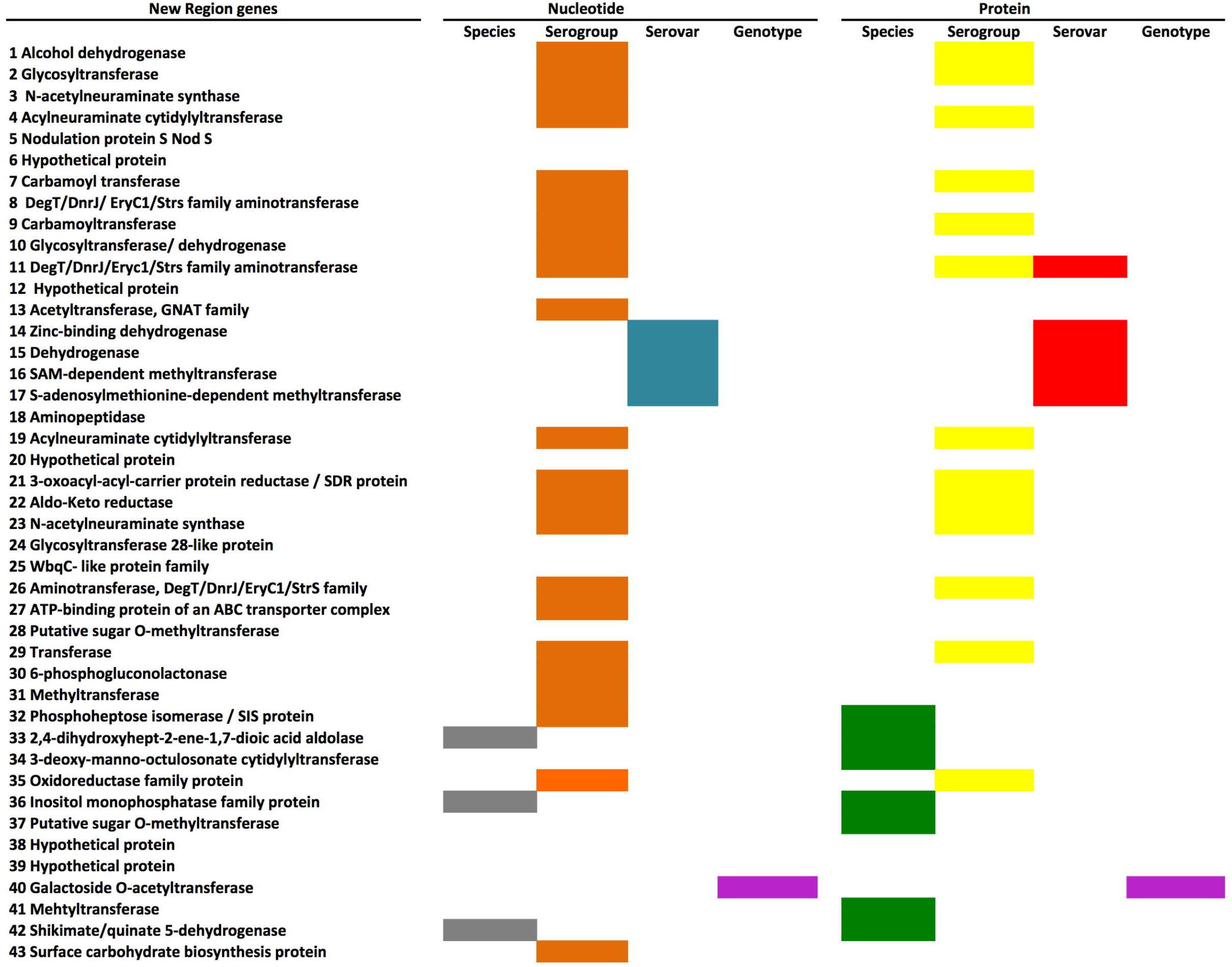
Correlation among Sru locus phylogeny and *Leptospira* spp. taxonomic classification. Heatmap of phylogenetic results of 43 nucleotide and protein predicted sequences. Colors diagram represents taxonomic Leptospira spp. degrees of classification. To nucleotide sequences, gray represents nucleotide sequences that clustering *L. interrogans* and *L. borgpertensenii* species, orange represents nucleotide sequences that clustering Seroj serogroup, blue represents nucleotide sequence that clustering Hardjo serovar, purple represents nucleotide sequence that clustering the Hardjoprajitno genotype. To protein sequences, green represents amino acid sequences that clustering *L. interrogans* and *L. borgpertensenii* species, yellow represents amino acid sequences that clustering Seroj serogroup, red represents amino acid sequences that clustering Hardjo serovar and purple represents amino acid sequences that clustering the Hardjoprajitno genotype.

The correlation of the *sru* locus within serological features was also evaluated in distinct *Leptospira* spp. that belonged to serovar Hardjo. The results obtained with *L. meyeri* serovar Hardjo showed that from a total of 43 genes there were 11 negative matches, 29 encoding genes with less than 40% identity, sequence coverage varying from 18% to 97%, and 3 genes showing identities higher than 40%. We also analyzed the predicted sequence of *L. borgpetersenii* serovar Ballum. The results showed a total of 27 proteins with less than 40% identity, negative results for 12 proteins and 4 genes with an identity higher than 40%.

Phylogenetic analysis using the protein sequence based on *sru* locus genes showed results that were similar to those of the nucleotide phylogenetic analysis. Thus, the detected divergences along the *sru* locus were maintained in the genomes, and they can affect protein activities, including different phenotypes in typing assays. A phylogenetic analysis based on protein sequences also demonstrated that the topology of most of the trees produced (14 gene trees) clustered the strains of *L. interrogans* and *L. borgpetersenii* of serogroup Sejroe together (Figure 5 and Supplementar Figures 1 and 2). Thus, the ortholog groups that produced phylogenetic trees in agreement with serogroup classifications were from the enzymes glycosyltransferase, acetylneuraminate cytidyltransferase, carbamoyltransferase, DegT aminotransferase, 3-oxoacyl reductase, N-acetylneuraminate synthase, putative sugar O-methyltransferase, transferase and oxidoreductase. These enzymes were previously associated with LPS and o-antigen cluster biosynthesis in *Escherichia coli*, *Legionella pneumophila*, *Pseudomonas aeruginosa, Vibrio cholerae, Campylobacter jejuni and L. interrogans* (22, 23, 24, 25, 26, 6). The surface antigens LPS and o-antigen are also targets of *Leptospira* serogroup Serjroe typing by MAT (1,, 27, 19), indicating that the identified predicted molecular patterns are in accordance with the current principal method for serogroup identification.

## Discussion

Leptospirosis remains an important zoonotic disease with a high impact in livestock production (28). Distinct serogroups and serovars of *Leptospira* spp. can result in different levels of disease severity, and serovars are directly correlated to specific hosts. In this regard, knowledge of the molecular divergence and evolution of *Leptospira* species can be useful for enhancing epidemiological disease control. In this study, we performed a comparative genomic analysis to identify molecular patterns that discriminate the *Leptospira* strains of serogroup Serjoe and serovar Hardjo. First, using whole-genome sequence analysis, we observed high similarities among *L. interrogans* serovars, however similar genome synteny and identities were observed among *L. interrogans* and *L. borgpetersenii* serovar Hardjo (13) (Supplementar Table 1 and Figure 2). These findings are in agreement with previously described results, where *L. interrogans* serovar Hardjo was found to possess identical synteny and identity of ORF15-ORF31 with that of *L. interrogans* serovar Copenhageni (6). However, in the current study, we observed that ORF15-ORF31 from *L. interrogans* possess higher identity with *L. interrogans* serovar Hebdomadis than with serovar Copenhageni. In fact, ORF22 in the *rfb* locus from *L. interrogans* serovar Hardjo possess high identity with *L. interrogans* serovar Hebdomadis str. C401 (19). Previous studies have described the Hebdomadis serogroup to be the largest and most geographically widespread of all the serogroups; it has consequently been divided into three autonomous serogroups–Hebdomadis, Mini and Sejroe (29).

In *Leptospira* spp., LPS and O-antigen biosynthesis is directly associated with the *rfb* locus (6). The whole-genome sequence analysis of *L. interrogans* serovar Copenhageni str. L1-130 suggested that the repetitive element in the *rfb* loci was likely acquired from lateral gene transfer (30, 31).

Previous work on the *L. interrogans* serovar Hardjo genotype Hardjoprajitno *rfb* locus found that ORF1 to ORF14 was probably acquired from *L. borgpetersenii* serovar Hardjo-bovis, and ORF15 to ORF31 were acquired from *L. interrogans* serovar Copenhageni L1-130 (11).

The *rfb* locus in *L. interrogans* serovar Hardjo possess a singular genetic organization that is correlated with the occurrence of a mobile element as part of the *rfb* locus in serovar Hardjo (6). IS3 mobile elements between ORF14 and ORF15 from the *rfb* locus were observed in *L. interrogans* str. Brem 329, *L. interrogans* serovar Hardjo str. Hardjo-prajitno and *L. interrogans* serovar Hardjo str. Norma. Based on sequence similarity and coverage, *L. interrogans* serovar Hardjo probably acquired ORF1-ORF14 from *L. borgpetersenii* serovar Hardjo-bovis, as previously hypothesized (11). Because ORF1 to ORF14 of the *rfb* locus from *L. interrogans* serovar Hardjo is very similar to that of *L. borgpetersenii* serovar Hardjo-bovis, we evaluated whether the genome section located upstream from the *rfb* locus showed the same pattern of conservation. This *sru* locus is composed of 43 genes and encompasses 45 Kb, with identical synteny within the *L. interrogans* and *L. borgpetersenii* strains that are classified in serogroup Sejroe, especially *L. borgpetersenii* serovar Hardjo-bovis. The *sru* locus also has a high synteny and sequence similarity with all strains classified in serogroup Sejroe, including *L. interrogans* serovar Hebdomadis, Medanensis, Hardjo str. Hardjo-prajitno and *L. borgpetersenii* serovar Hardjo-bovis, suggesting genetic acquisition or genetic replacement. Moreover, the occurrence of an insertion sequence downstream of the *sru* locus located between ORF14 and ORF15 in the *rfb* locus in *L. interrogans* serovar Hardjo and *L. borgpetersenii* serovar Hardjo corroborates the hypothesis of genome recombination in this site.

Genes in the *sru* locus from the *L. interrogans* serovar Hardjo genome were predicted to encode enzymes such as glycosyltransferase, acetylneuraminate cytidyltransferase, carbamoyltransferase, DegT aminotransferase, 3-oxoacyl reductase, N-acetylneuraminate synthase, putative sugar O-methyltransferase, transferase and oxidoreductase. All of these enzymes are associated with carbohydrate biosynthesis (22,23, 24, 25, 26, 6) Cote et al., 2017, Klein et al., 2009, Farhat et al., 2011, King et al., 2009, Aydanian et al., 2011, Kalynych et al., 2014, Kalambaheti et al., 1999) and are predicted to be involved specifically with LPS and O-antigen biosynthesis in *Escherichia coli*, *Legionella pneumophila*, *Pseudomonas aeruginosa, Vibrio cholerae, Campylobacter jejuni* and *L. interrogans* (22, 23, 24, 25, 26, 6). In addition, for several bacteria species, LPS is directly associated with pathogenic processes and disease development (32). In *Leptospira* spp., LPS is involved in several processes and is associated with diagnosis, immune response and taxonomic classification (1, 6, 33, 18). In leptospirosis diagnosis, the surface antigens LPS and o-antigen are targets of the *Leptospira* serogroup Sejroe typing by the seroagglutination test (MAT), indicating that the predicted molecular patterns identified are in accordance with the current principal method for serogroup identification and that the key concept is seroconversion on paired samples associated with LPS/*rfb* locus leptospiral composition (1,6,11). Thus, the detected divergences along the *sru* locus were fixed in the genome; they can affect protein activities including different phenotypes in typing assays.

The phylogenetic analysis of nucleotide and amino acid sequences from *sru* locus genes revealed that most of the genes clustered strains from serogroup Sejroe together. Among all *sru* locus ORFs, aminotransferase protein may be relevant for the molecular typing of *Leptospira* spp. because the sequence is present in all *Leptospira* strains and it clustered all strains of serogroup Sejroe in the analysis. Related literature has discussed the application of aminotransferase proteins in the phylogenetic analysis of bacteria. In *Halomonas variabilis*, aminotransferase phylogeny results clustered strains from different geographic locations together (Okamoto et al., 2008). Here, we observed that aminotransferase phylogeny also showed a topology that clustered strains from the same geographic locations, such as strains from Brazil, Indonesia and Germany. Based on this *in silico* analysis, the next step in this work is to evaluate *Leptospira* spp. encoding region genotypes by applying genetic methodologies.

## Conclusion

This study suggests that the recombination site identified in the *L. interrogans* serogroup Sejroe serovar Hardjo subtype Hardjoprajitno genome is composed of sugar metabolism genes that could be directly associated with *Leptospira* spp. serogroup classification. This region is also conserved within serogroup Sejroe. Including *sru* locus genes in studies of the *rfb* locus could improve the molecular epidemiology analysis and understanding of *Leptospira* spp. diversity.

## Supporting information

Proteome comparison analysis among L. interrogans and L. borgpetersenii species.

The BLAST identity of rfb nucleotide locus from Leptospira spp.

The BLAST identity of Sru nucleotide locus from Leptospira spp.

Gene ontology analysis of Sru locus from L. interrogans serovar Hardjo.

Leptospira strains used in phylogenetic analysis of Sru locus.

Phylogenetic trees using nucleotide and protein sequences of Sru locus in Leptospira spp.

## Funding Informations

This study was supported by Conselho Nacional de Desenvolvimento Científico e Tecnológico (CNPq) and Instituto Nacional de Ciência e Tecnologia (INCT) de Informação Genético-Sanitária da Pecuária Brasileira.

## Author Contributions

Conceptualization, M.R.V.C., T.A.O.M., J.M.O. and T.S.; methodology, M.R.V.C. and T.A.O.M.; validation M.R.V.C., T.A.O.M. and T.S..; formal analysis, M.R.V.C., T.A.O.M. and T.S., investigation, M.R.V.C., T.A.O.M. and T.S..; resources, R.C.L., J.P.A.H., E.C.M., T.A.O.M.,.; writing – original draft preparation, M.R.V.C., T.A.O.M.and T.S.; writing – review and editing, M.R.V.C., T.A.O.M and T.S.; visualization, M.R.V.C, T.A.O.M., T.S.; supervision, T.A.O.M., J.M.O..; project administration, T.A.O.M., J.P.A.H. and J.M.O.; funding, R.C.L., E.C.M, J.P.A.H.

## Acknowledgments

The authors would like to thank Conselho Nacional de Desenvolvimento Científico e Tecnológico (CNPq) and Instituto Nacional de Ciência e Tecnologia (INCT) de Informação Genético-Sanitária da Pecuária Brasileira and FAPEMIG.

---

**Supplementar Table 1: Proteome comparison analysis among *L. interrogans* and *L. borgpetersenii* species.** Comparative analysis of whole genome sequence from *L. interrogans* serovars Hardjo str. Norma, str. Hardjo-prajitno and str. Brem 329, serovar Copenhageni str. L1-130, serovar Lai str. 56601 and *L. borgpetersenii* serovar Hardjo-bovis str. L550.

**Supplementar Table 2: The BLAST identity of *rfb* nucleotide locus from *Leptospira* spp.**

**Supplementar Table 3: The BLAST identity of *Sru* nucleotide locus from *Leptospira* spp.**

**Supplementar Table 4: Gene ontology analysis of *Sru* locus from *L. interrogans* serovar Hardjo.**

**Supplementar Table 5: *Leptospira* strains used in phylogenetic analysis of *Sru* locus.**

**Supplementar Figure 1: Phylogenetic trees using nucleotide sequences of *Sru* locus in *Leptospira* spp**. The nucleotide sequences were aligned using Muscle, and the trees were generated using the maximum likelihood method with model maximum composite likelihood and 1,000 bootstrap. Gray and orange represent clusters of *Leptospira* species and serogroup Sejroe, respectively.

**Supplementar Figure 2: Phylogenetic trees using protein sequences of *Sru* locus in *Leptospira* spp**. The protein sequences were aligned using Muscle, and the trees were generated using the maximum likelihood method with model maximum composite likelihood and 1,000 bootstrap. Green, yellow and red branches represent clusters of *Leptospira* species, serogroup Sejroe and serovar Hardjo, respectively.

