## Supplementary material for "A potential genomic recombination site upstream of the *rfb* locus in *Leptospira interrogans* is associated with serogroup Serjoe and serovar Hardjo classification": Phylogenetic trees using nucleotide and protein sequences of Sru locus in Leptospira spp.

A) Carbamoyltrasnferase

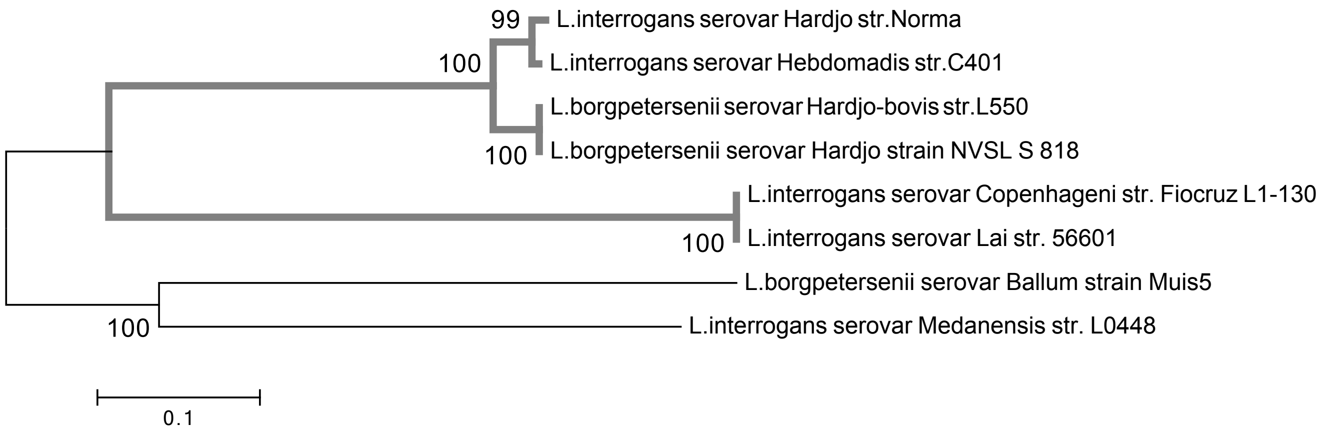


B) Deg1 Aminotransferase

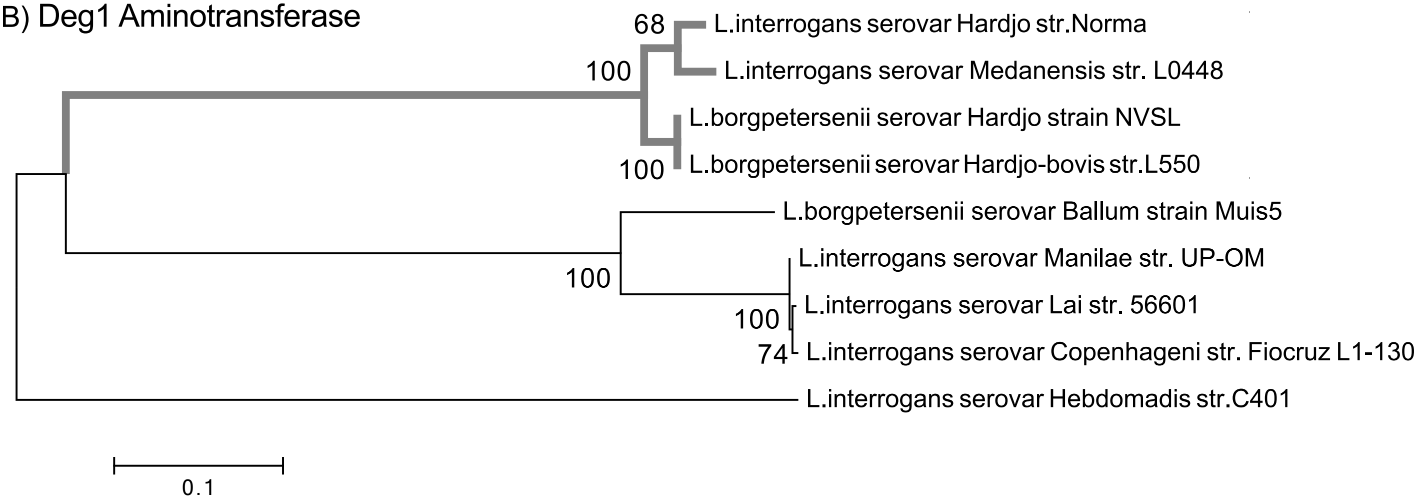


C) Carbamoyltransferase

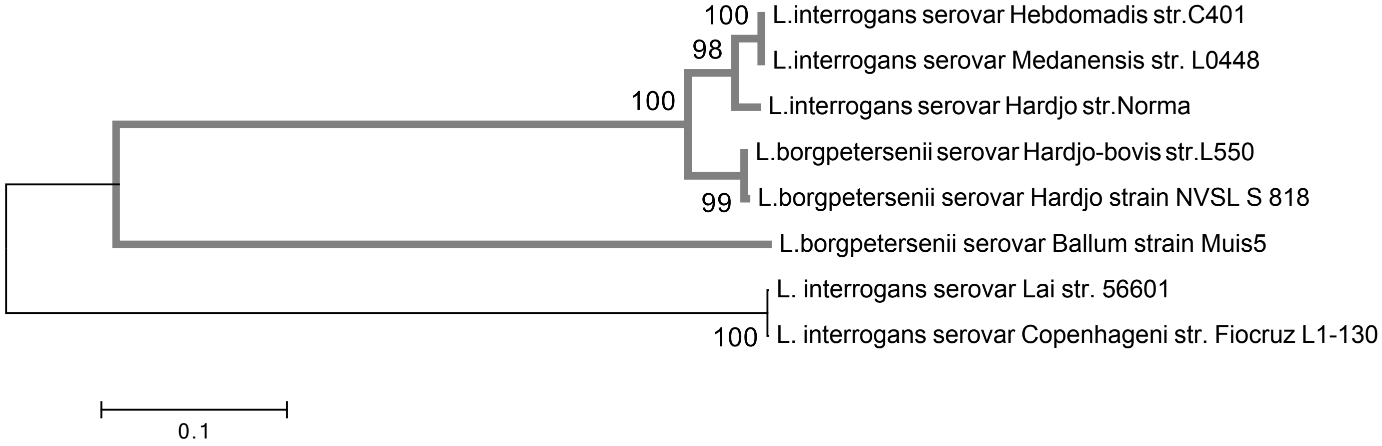

D) Aminopeptidase

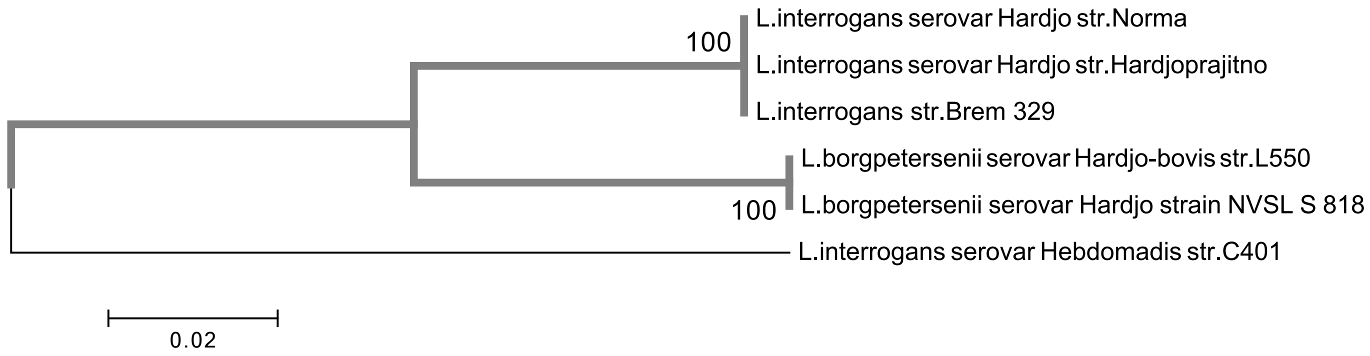


E) 6-Phosphogluconolactonase

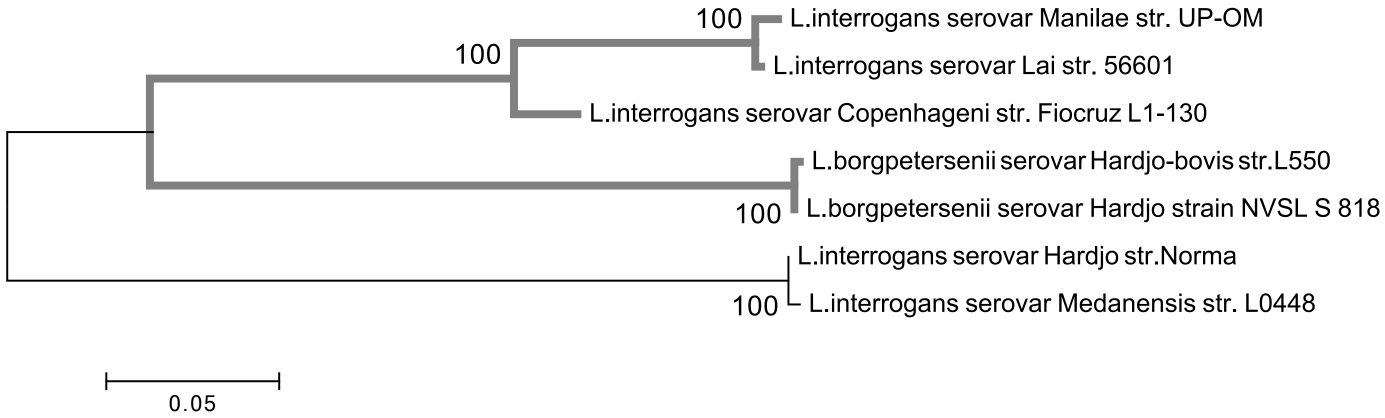


F) 2,4 dihydroxyhept-2-ene-1,7-dioic acid aldolase

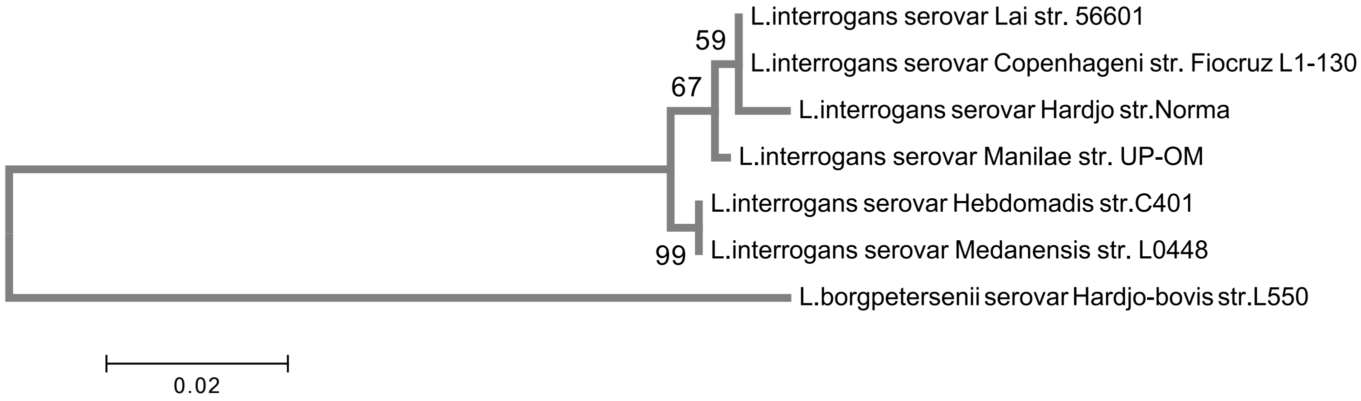

G) Inositol Monophosphatase

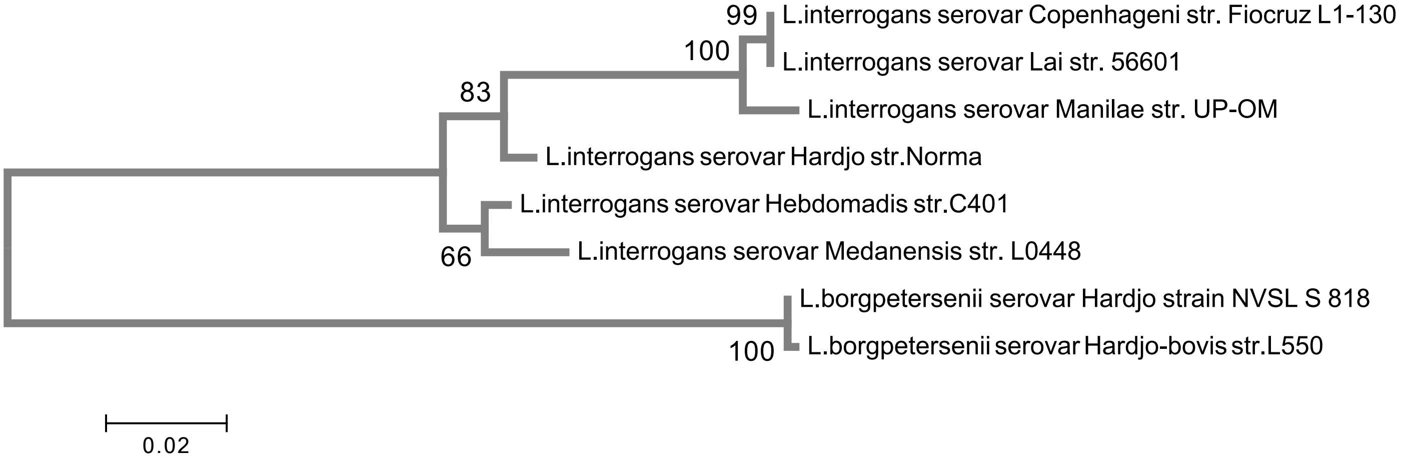


H) Putative Sugar O-Mehtyltransferase

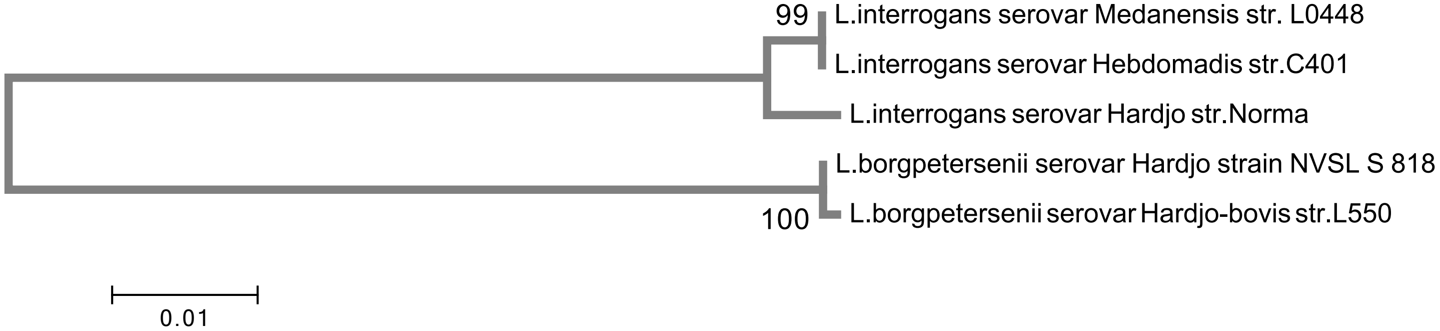


I) Mehtyltransferase

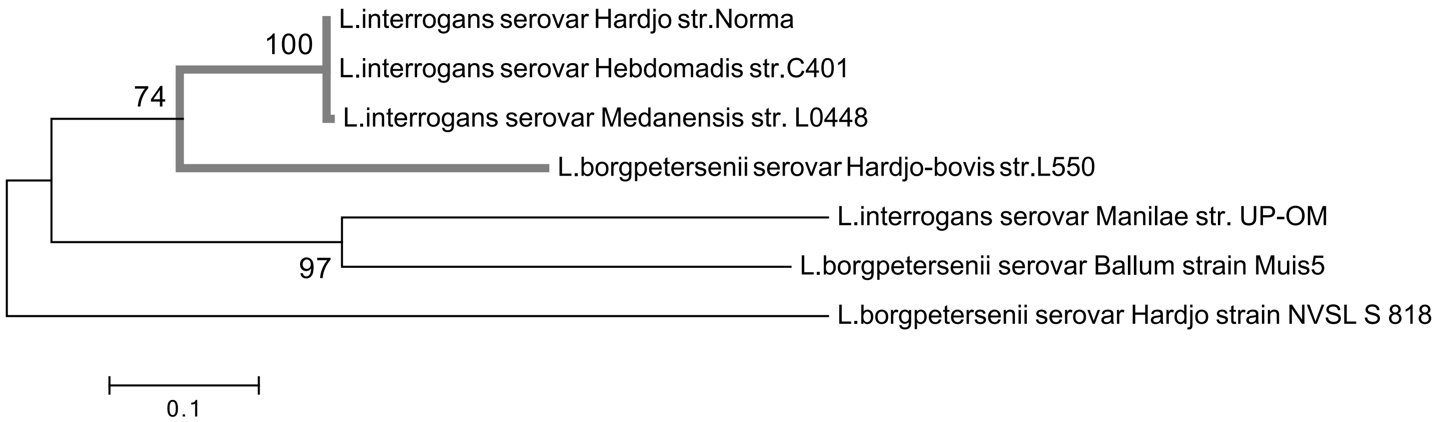


J) Shikimate/ Dehydrogenase

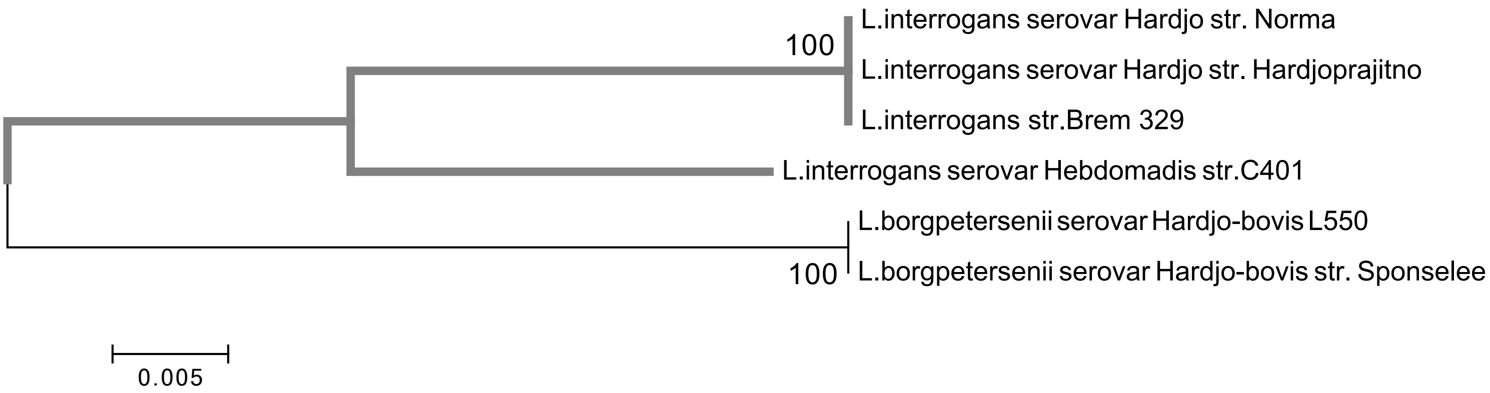
